# Shaping liposomes by cell-free expressed bacterial microtubules

**DOI:** 10.1101/2021.06.13.448053

**Authors:** Johannes Kattan, Anne Doerr, Marileen Dogterom, Christophe Danelon

## Abstract

Genetic control over a cytoskeletal network inside lipid vesicles offers a potential route to controlled shape changes and DNA segregation in synthetic cell biology. Bacterial microtubules (bMTs) are protein filaments found in bacteria of the genus *Prosthecobacter*. They are formed by the tubulins BtubA and BtubB which polymerize in the presence of GTP. Here, we show that the tubulins BtubA/B can be functionally expressed from DNA templates in a reconstituted transcription-translation system, thus providing a cytosol-like environment to study their biochemical and biophysical properties. We found that bMTs spontaneously interact with lipid membranes and display treadmilling. When compartmentalized inside liposomes, de novo synthesized BtubA/B tubulins self-organize into cytoskeletal structures of different morphologies. Moreover, bMTs can exert a pushing force on the membrane and deform liposomes, a phenomenon that can be reversed by light-activated disassembly of the filaments. Our work establishes bMTs as a new building block in synthetic biology. In the context of creating a synthetic cell, bMTs could help shape the lipid compartment, establish polarity or directional transport, and assist the division machinery.

## INTRODUCTION

In eukaryotic cells, microtubules formed by the polymerization of α- and β-tubulin proteins are involved in essential functions, such as intracellular transport and chromosome segregation. Bacterial tubulin homologues have recently been discovered in *Prosthecobacter* species [1,2]. Called bacterial tubulin A and B (BtubA/B), these proteins interact to form microtubule-like structures in the presence of GTP [3]. Bacterial microtubules (bMTs) consist of five [4] or four protofilaments [5], as reported in in vivo and in vitro studies, respectively. Bacterial microtubules are thus noticeably thinner than eukaryotic microtubules which consist of 13 protofilaments. Recent in vitro studies have shown that bMTs exhibit dynamic instability (stochastic switching between growth and shrinkage) and treadmilling (apparent directional movement caused by the net addition of new subunits on one end and net removal on the other) [5,6]. These properties are also common to eukaryotic microtubules.

The function of bMTs remains unclear. *Prosthecobacter* itself belongs to the Phylum *Verrucomicrobia* and consists of Gram-negative bacteria which exhibit a high degree of compartmentalization. *Prosthecobacter dejongeii*, for example, possess a major membrane-bounded region, containing the fibrillar nucleoid and all the ribosome-like particles, as well as an intracytoplasmic membrane [7]. Another distinguishing feature of *Prosthecobacter* is the presence of narrowed extensions of the cell wall, called prosthecae [4]. Bacterial microtubules seem to be predominately located in these cell stalks, which suggests that they might be involved in their formation. It has also been proposed that BtubA/B filaments may contribute to intracellular organization [8]. The only currently known protein that interacts with bMTs is BtubC (also known as bacterial kinesin light chain, Bklc) which stabilizes bMTs [5] and links them to lipid membranes in vitro [8], suggesting it could play a role in anchoring BtubA/B filaments to membrane protrusions in vivo.

Unlike eukaryotic tubulin, BtubA/B is not dependent on protein chaperones and post-translational modifications, and can be functionally expressed in *E. coli* [4]. Because the cytoplasm of *Prosthecobacter* probably resembles that of *E. coli* [3], cell-free protein synthesis (CFPS) platforms derived from *E. coli* represent a physiologically relevant environment to investigate bMTs. Not only does CFPS allow to bypass protein purification, but it also enables the continuous interrogation of the protein dynamic behaviour in the course of its production [9].

To better apprehend the properties of bMTs in a cytosol-like environment, while taking advantage of the versatility of cell-free assays, we reconstituted in this study BtubA/B in a CFPS system. To further mimic the membrane-rich environment in *Prosthecobacter*, we characterized cell-free expressed BtubA/B on supported phospholipid bilayers and within vesicle compartments. Our results demonstrate that active BtubA/B proteins can be produced from their genes in vitro. Moreover, de novo synthesized bMTs can assemble on supported membranes and inside lipid vesicles (even without BtubC), where they form cytoskeletal structures that can deform the vesicle membrane. As CFPS inside liposomes has become an attractive platform to build a synthetic cell [10-17], we believe that bMTs could be exploited for spatial organization, polarization, and shape transformation of artificial cell models.

## RESULTS AND DISCUSSION

We chose the PURE system [18], a reconstituted *E. coli*-based translation machinery (specifically the commercially available PURE*frex*2.0), as our CFPS platform. This choice was motivated by the very low levels of protease and nuclease activity, and the wide range of (membrane) proteins synthesized in an active state with this system [12-20]. DNA templates for PURE*frex*2.0 reactions consisted of the *btubA* and *btubB* genes from *Prosthecobacter dejongeii*. Both DNA constructs were sequence optimized for (i) expression in an *E. coli* host by matching codon occurrence with tRNA usage, (ii) low GC content within the first 30 base pairs (synonymous mutations were introduced to keep the amino acid sequence unaltered), and (iii) low propensity of intramolecular base-pairing of the messenger RNA around (in the vicinity or involving) the start codon (**Supplementary Fig. 1**). To satisfy the latter condition, the change in free Gibbs energy (Δ*G*) for a few RNA molecules was calculated. Lower Δ*G* values represent higher melting temperatures of the RNA molecule, which is known to be potentially inhibitory of translation in the PURE system [21]. Thus, we selected for each construct the DNA sequence whose corresponding mRNA has the predicted lowest melting temperature, assuming this would decrease the occurrence of inhibitory secondary structures involving the ribosome binding site and the start codon.

Proteins BtubA/B (**Fig. 1A**) were first separately expressed from linear DNA templates in PURE*frex*2.0, supplemented with FluoroTect GreenLys reagent to fluorescently label gene products through co-translational incorporation of lysine residues conjugated to a fluorophore (**Fig. 1B**). Analysis of samples by denaturing polyacrylamide gel electrophoresis (PAGE) confirmed in vitro production of each Btub protein (**Fig. 1C, Supplementary Fig. 2**). The concentration of expressed BtubA/B was estimated by using a standard curve with purified proteins where values of ∼13 μM (BtubA) and ∼17 μM (BtubB) were obtained (**Supplementary Fig. 3**). Subsequent co-expression from an equimolar amount of the *btubA* and *btubB* genes yielded ∼3 μM of BtubA and ∼5 μM of BtubB (**Supplementary Fig. 3**). It is unclear why the total concentration of synthesized proteins is lower in co-expression assays compared to separate expression of single genes.

**Figure 1:**
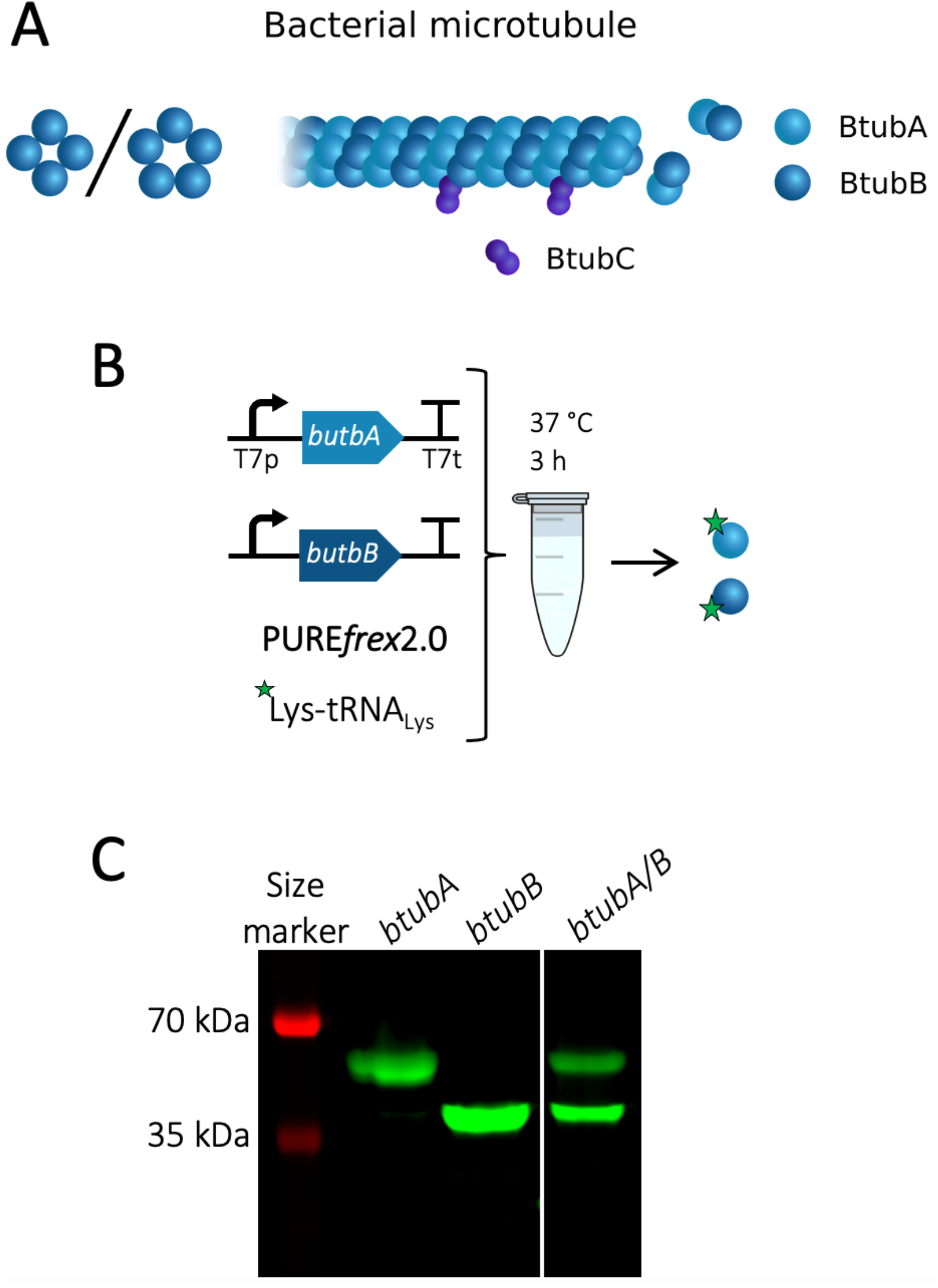
Cell-free expression of BtubA/B. (A) Schematic of a bacterial microtubule consisting of the BtubA/B subunits arranged in four to five protofilaments. The protein BtubC binds to the outside of the filament, predominantly interacting with BtubB. (B) Schematic of CFPS reaction. PURE*frex*2.0 was supplemented with the GreenLys reagent to fluorescently label the synthesized proteins through BODIPY-conjugated lysine residues. (C) SDS-PAGE analysis of the gene expression products. The gene names of the expressed DNA templates are indicated on the lanes. The uncropped gel is shown in **Supplementary Fig. 2**.

Next, we examined the activity of purified and cell-free synthesized bMTs on supported lipid bilayers (SLBs). The membrane composition in SLB assays consisted of ∼50 mol% of phosphatidylethanolamine (PE) and phosphatidylglycerol (PG) lipids, which are also found in the membrane of *Prosthecobacter* [22]. Purified BtubA/B proteins recombinantly expressed in *E. coli* cells were investigated in a PURE*frex*2.0 background to closely emulate the molecular and ionic complexity of the bacterial cytosol. Bacterial microtubules were successfully assembled on top of an SLB from 2.5 μM purified BtubA/B doped with Atto488-labelled subunits for visualization. The bMTs localized exclusively on the SLB and not on the bare glass areas (**Fig. 2A**). Filaments of BtubA/B appeared to stably interact with the membrane without the need for anchoring proteins such as BtubC. The protein BtubC synthesized in PURE*frex*2.0 was able to bind to lipid membranes and to promote recruitment of BtubA/B (**Supplementary Fig. 4 and Supplementary Note**). Moreover, bMTs formed bundles of multiple protofilaments over time, suggesting lateral interactions. The critical BtubA/B concentration for the assembly of bMTs was reported to be 2.5-5 µM at the potassium concentration present in the PURE system [6]. Here, bMTs were observed on SLBs already at concentrations of ∼1 µM of purified tubulin in PURE*frex*2.0 (**Supplementary Fig. 5**). This result suggests that, in the arguably more physiological conditions used here (in particular the higher molecular crowding), the critical concentration for BtubA/B polymerization is lower. Differences due to the types of activity assays (high-speed pelleting of BtubA/B and filaments grown from seeds in [6]) may also explain this change.

**Figure 2:**
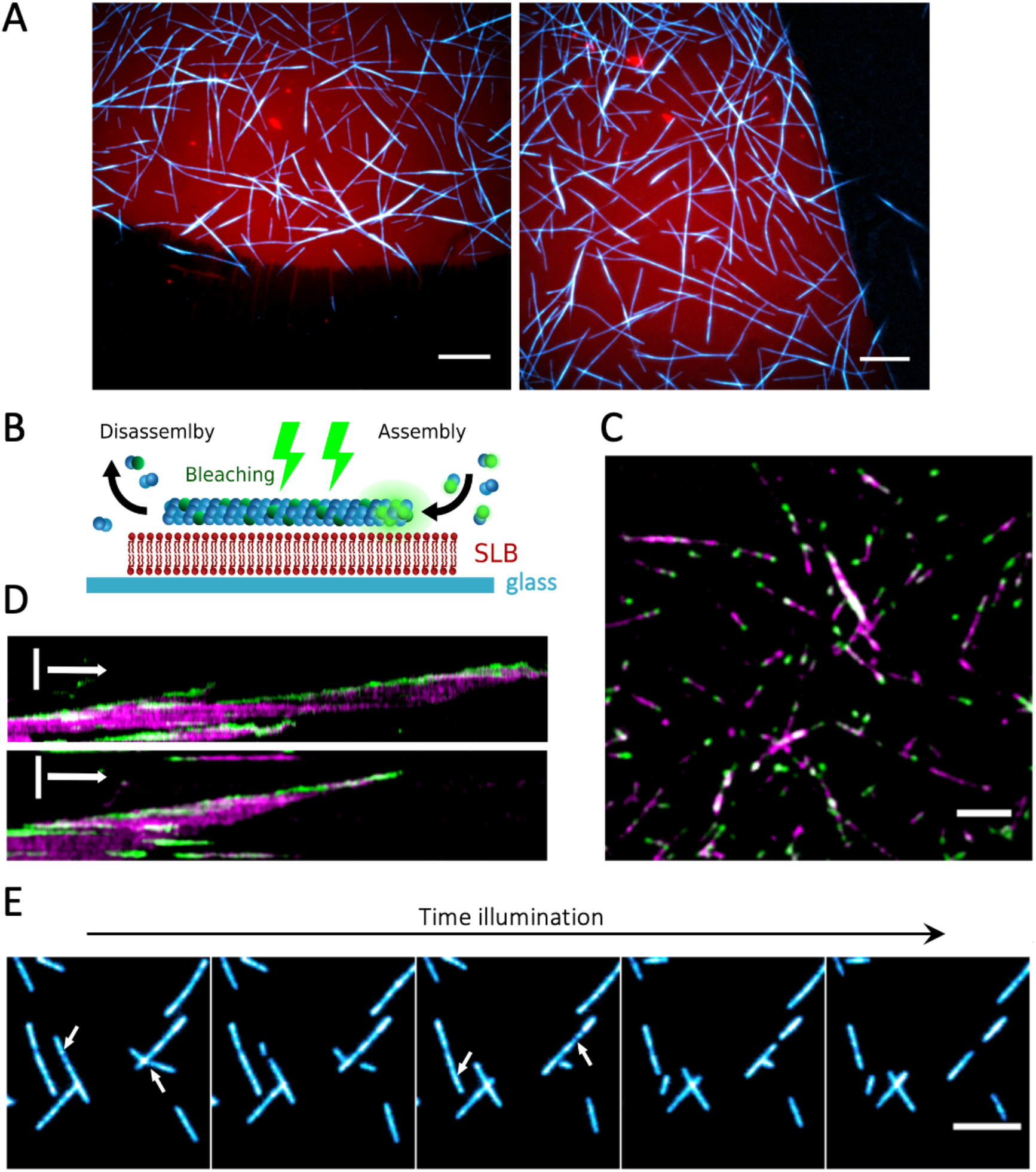
Dynamics of bMTs formed by purified proteins on SLBs. (A) Fluorescence microscopy images showing selective binding of bMTs (cyan, Atto488-BtubA/B) onto an SLB (red, DHPE-TexasRed). Concentration of bacterial tubulins was 2.5 μM. Scale bars: 10 µm. (B) Schematic of the dual colour labelling assay to study bMT dynamics on an SLB. Purified BtubA/B proteins labelled with either Atto488 or Atto565 were used. Atto488-labelled bacterial tubulins were bleached continuously during imaging. The only active Atto488 fluorophores are located at the plus end of the filament, where new subunits incorporate. (C) Bleached filaments display fluorescent ends originating from continuous addition of fresh, unbleached Atto488-BtubA/B (green). Atto565-BtubA/B is coloured in magenta. Scale bar: 5 µm. (D) Kymographs showing bMT dynamics during photobleaching. Vertical scale bar: 5 µm. Horizontal arrow: 1 min. (E) Time series images showing bMT disassembly events during high-intensity illumination. Arrows indicate the locations at which the filaments break apart. Duration was 120 s between the first and last image. Scale bar: 5 µm.

Single filaments appear to undergo directional movement along their longitudinal axis (**Movie 1**). This behaviour is presumably caused by simultaneous growth and shrinkage on the two opposite ends of the filament, a phenomenon known as treadmilling, as was previously also observed for purified BtubA/B filaments in simple buffers [5,6]. To validate this hypothesis, we used dual colour labelling of tubulin with the Atto488 or Atto565 dye, and bleached one of the fluorophores (Atto488) during imaging (**Fig. 2B**). Continuous illumination yielded filaments displaying extensions with fluorescent extremities originating from addition of tubulin subunits to the growing ends (**Fig. 2C, D, Movie 2**). Further, single fluorescent spots on the bleached filaments remained immobile. These observations demonstrate that bMTs undergo treadmilling behaviour on SLBs in a solution compatible with CFPS.

When cell-free expressed BtubA/B proteins, along with a trace amount of labelled purified bacterial tubulin (100 nM), were added to an SLB (**Fig. 3A**), assembly of bMTs and recruitment to the SLB were immediately observed (**Fig. 3B**). To rule out the possibility that filament assembly might be caused by the low amount of labelled purified tubulin, we also imaged the SLB with expressed BtubA and 100 nM of labelled tubulin for 30 min. No bMTs were observed in the absence of expressed BtubB, but readily appeared again upon addition of expressed BtubB (**Fig. 3C, Movie 3**). As with the purified proteins, dynamic instability, treadmilling on the membrane, as well as bundling of the microtubules were observed (**Movie 3**).

**Figure 3:**
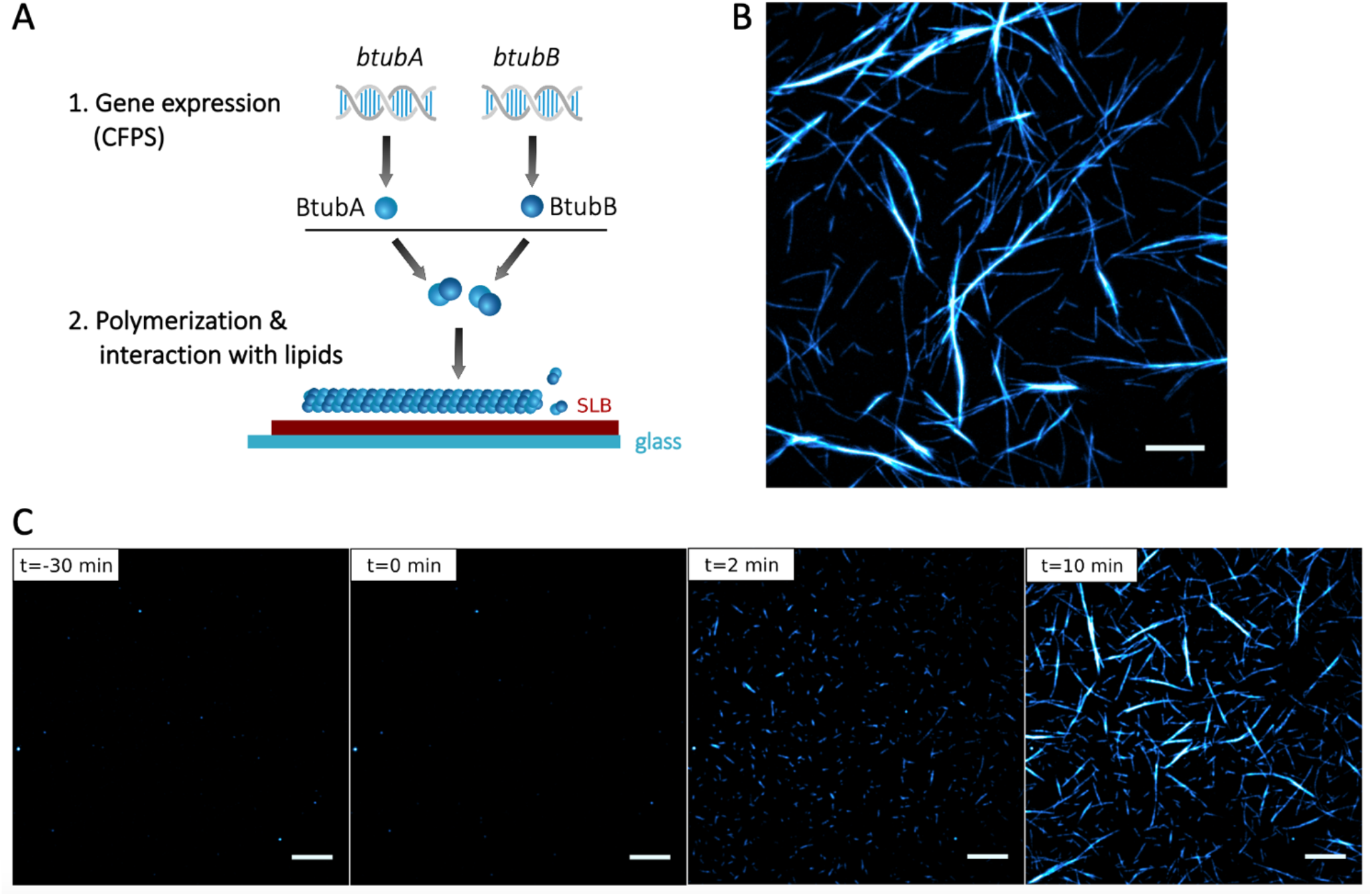
Cell-free expressed BtubA/B assemble into dynamic bMTs on SLBs. (A) Schematic of CFPS and BtubA/B polymerization on supported lipid membranes. (B) Fluorescence microscopy image of cell-free expressed bMTs containing a small fraction of purified Atto488-BtubA/B. Scale bar: 10 µm. (C) Cell-free synthesized BtubA was mixed with 100 nM Atto488-BtubA/B and incubated on an SLB for 30 min. No filament was observed. At time *t*=0, separately expressed BtubB was added on top of the SLB, triggering immediate formation of short filaments that developed into longer and thicker bundles. Scale bars: 10 µm.

When the bMTs on the SLB were imaged at high illumination intensity, the filaments could break apart, either completely disassembling or with generated fragments detaching from the membrane (**Fig. 2E**). This effect was dependent on the presence of the fluorophore used for labelling (Atto488 or Atto565), suggesting a dye-specific photochemical reaction. If excitation was performed using a laser with a wavelength outside the excitation range of the fluorophore, prolonged illumination was not accompanied with filament breaking. Dual labelling experiments confirmed that the physical integrity of bMTs was altered, as opposed to photobleaching effects (**Movie 4**).

Compartmentalization of bMTs in cell-sized lipid vesicles provides a unique platform to study their self-organization in a closed volume, as well as their ability to exert pushing forces and deform the membrane. Large and giant liposomes were produced by glass bead-assisted lipid film swelling [23,24]. PURE*frex*2.0, the *btubA/B* DNA constructs, a mix of DnaK chaperone to promote protein folding [9], and 100 nM of Atto488-labelled purified BtubA/B were co-encapsulated (**Fig. 4A**). A slightly higher concentration of the *btubA* gene compared to the *btubB* gene (3.75 nM vs 2.5 nM) was used to compensate for the lower amount of expressed BtubA vs BtubB when an equimolar concentration of the two templates is used (**Fig. 1C**). Liposomes that sedimented in a glass chamber were incubated at 37 °C to trigger gene expression and imaged at various time points by fluorescence confocal microscopy. Initially, liposomes that had encapsulated the labelled tubulin displayed an even fluorescence in their lumen (**Supplementary Fig. 6**). During the course of gene expression, filament structures appeared, with length and bundling propensity increasing over time (**Fig. 4B, Supplementary Fig. 6**). This internal cytoskeleton could clearly be attributed to in situ synthesis of BtubA/B as the small fraction of purified labelled tubulin did not yield filaments until accumulation of sufficient expression products (**Supplementary Fig. 6**) nor in control samples where the *btubA/B* genes were omitted (**Supplementary Fig. 7**). The number of liposomes exhibiting cytoskeletal structures, the time of incubation until the first filaments appear (typically 1 h), and the number of filaments per liposome, differed from one experiment to the other. Yet, the emergence of bMT networks was a robust observation. In the absence of DnaK mix, synthesized BtubA/B could not develop into cytoskeletal structures, suggesting that chaperones are needed to prevent synthesized proteins from aggregating or misfolding in liposomes [25]. A similar observation was made when expressing active Min proteins in PURE*frex*2.0 [9]. A variety of BtubA/B filament and liposome morphologies was seen: straight or curved bundles or meshes, located across the lumen or near the membrane (**Fig. 4C**). Most liposomes containing bMTs exhibited morphological changes. Vesicles that were originally spherical showed local protrusions or global elongation (**Fig. 4B, C**). Such deformation events were likely the result of the pushing force exerted by growing bMTs on the liposome membrane, a phenomenon that is well documented for eukaryotic microtubules and actin filaments [26,27]. Similar results were obtained upon direct encapsulation of purified BtubA/B (6.6 µM final concentration in PURE*frex*2.0), with the only noticeable difference that the average occurrence of filaments was higher with purified proteins, and a few cross-shaped liposomes were observed (**Fig. 5**). Liposome-to-liposome heterogeneity regarding the internal concentration of labelled tubulin, expression efficiency, and cytoskeleton features were likely the consequence of varying encapsulation efficiency and stochastic effects, as previously reported for other reconstituted biological systems [9,12,16,17,23,24].

**Figure 4:**
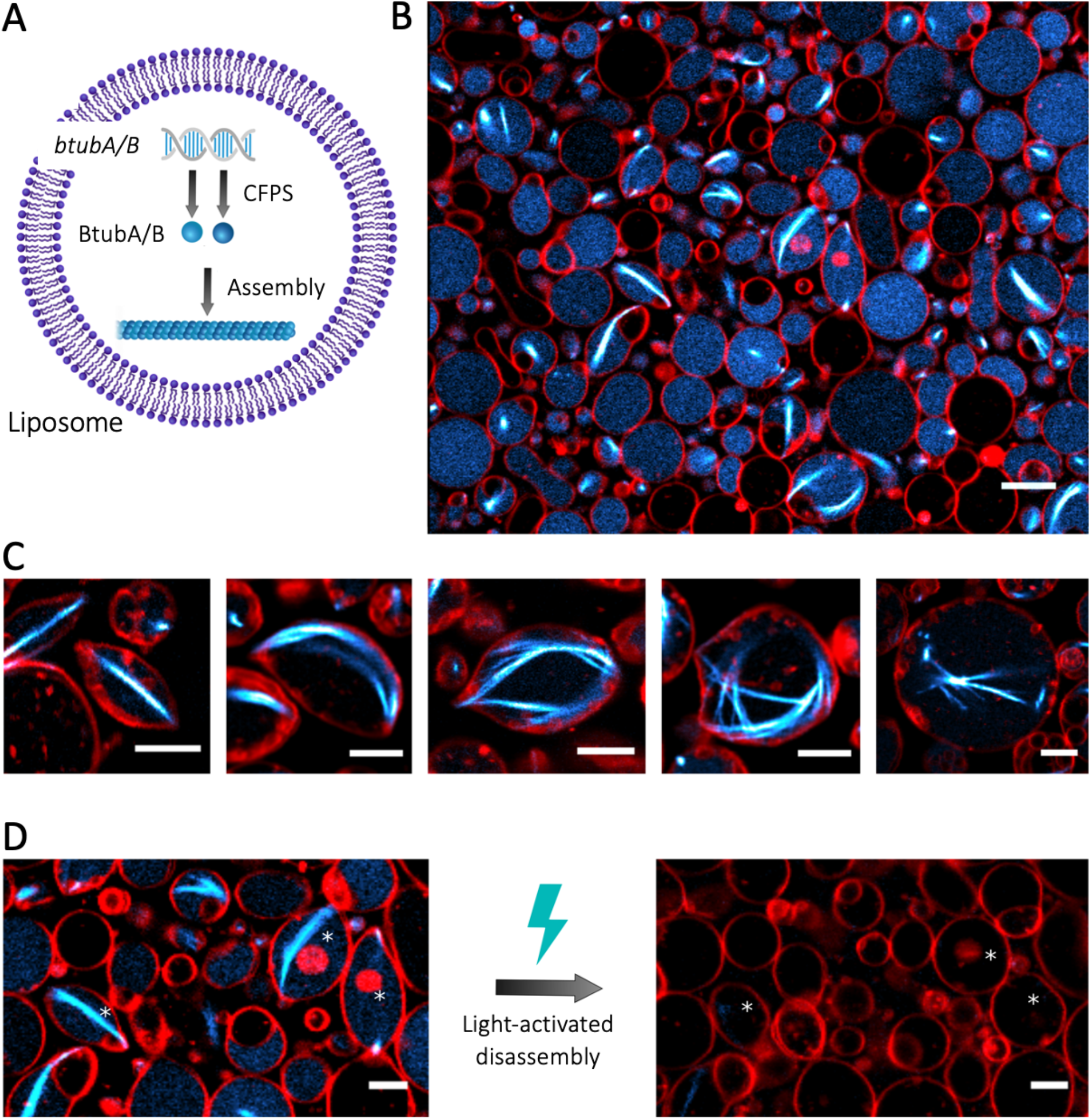
CFPS and assembly of bMTs inside liposomes. (A) Schematic of liposome-compartmentalized gene expression and synthesis of BtubA/B that self-organize into bMTs. (B) Fluorescence confocal microscopy images of liposomes (red, DHPE-TexasRed) with encapsulated Atto488-BtubA/B (100 nM, cyan) after 4 h CFPS reaction. In situ synthesized bMTs (cyan) are visible in several liposomes. Scale bar: 10 µM. (C) Examples of different bMT cytoskeletal structures showing clear membrane deformation. Samples were observed after 5 h of incubation at 37 °C. Scale bars: 5 µm. (D) Breaking of bMTs and relaxation of liposomes into a spherical shape was triggered by exposing samples to high laser intensity. Asterisks indicate liposomes whose shape was modified by light-activated disassembly of bMTs. Scale bars: 5 µm.

**Figure 5:**
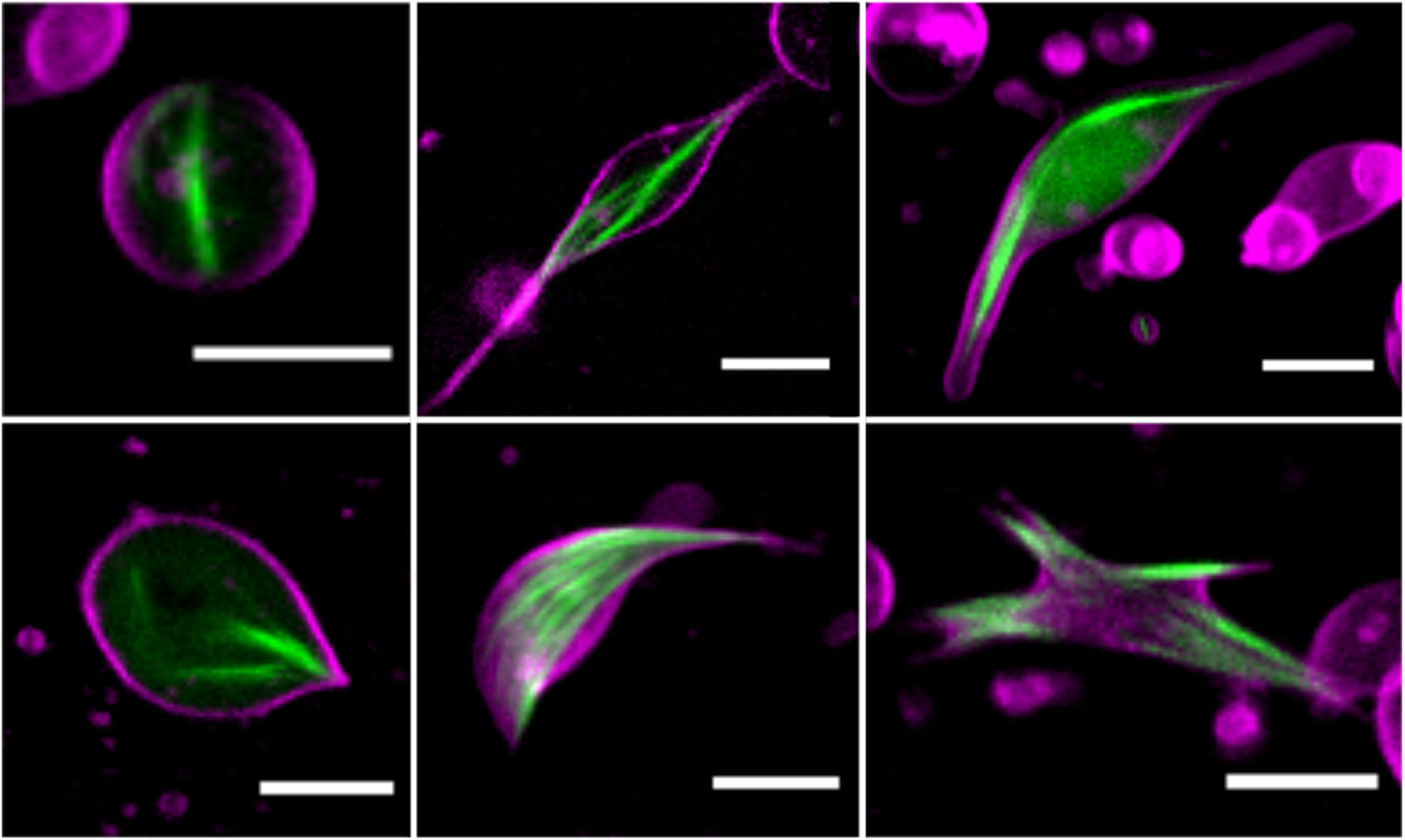
Bacterial microtubules (green, 100 nM Atto488-BtubA/B) formed by 6.6 μM of purified tubulins inside liposomes (magenta). Scale bars: 5 μm.

Finally, we sought to demonstrate the reversible nature of liposome deformations by the light-activated bMT disassembly mechanism that we observed on SLBs (**Fig. 2E**). Exposing filament-containing liposomes to intense illumination triggered bMT collapse and subsequent vesicle relaxation to a spherical shape (**Fig. 4D**). Similar observations have been reported with actin filaments [28]. In some cases, sphericality was not fully restored, which might be caused by high packing of liposomes or by surface tethering over an extended area, both effects altering the mechanical properties of the vesicles. Overall, these results demonstrate that liposome-confined, de novo synthesized bMTs can self-organize into cytoskeletal structures that can deform the liposome through pushing forces on the membrane. Moreover, light-activated bMT disassembly can be exploited for reversible shape control.

To date, quantitative insights about the physical parameters describing bMT mechanical properties are missing. The bending rigidity of bMTs is probably lower than that of eukaryotic microtubules due to their smaller diameter [4,5]. The BtubA/B filament and bundle morphologies share similarities with those reported with encapsulated actin filaments bundled by linker proteins [29]. Therefore, we assume that the rigidity of bMTs lies in the range between eukaryotic microtubules and actin filaments. Other relevant parameters that remain to be determined include the critical force for buckling, the pushing force of a growing bMT, and the number of filaments per bundle. This information will help design liposome experiments, where membrane tension and BtubA/B expression levels may be adjusted to modulate the vesicle shape and organization of cytoskeletal structures.

This work expands the scope of bMT applications in synthetic biology. Bacterial microtubules have structural and biochemical properties that give them decisive advantages for bioengineering compared to eukaryotic microtubules. Their comparatively lower threshold of monomer concentration for polymerization, ability to form bundles without additional cofactors, and suitability for cell-free gene expression represent assets for implementation in artificial cells. Other prokaryotic protein filaments have already been reconstituted in liposomes. MreB expressed in liposomes assembles into cytoskeletal networks without inducing membrane deformation [30], unless molecular crowding at the vesicle membrane is applied [31]. Rings of FtsZ-FtsA can form budding necks and constrict liposomes [17]. Here, we show that bMTs expand the repertoire of protein filaments and range of associated functions. Endowing liposomes with a DNA-programmed mechanism for elongation and membrane remodelling, as realized here with bMTs, might be instrumental to support processes such as polarization and compartment division. For instance, bMT-mediated elongation of vesicles may promote pole-to-pole oscillations of the reconstituted Min system [9], or the positioning of the Z-ring [17]. The prospect for discovering new bMT-interacting partners (natural or engineered), especially end-tracking proteins, could further extend their role in intravesicular transport, segregation of synthetic chromosomes, or polarization.

## MATERIALS AND METHODS

### Preparation of DNA constructs

To ensure a high expression level, we adjusted the DNA sequence of the three *btub* genes of *Prosthecobacter dejongeii* (*btubA, btubB, btubC*) with respect to their GC content and intramolecular base pairing for the first 30 bp after the start codon. For each gene, we listed all possible DNA sequences of the first 30 bp encoding the same amino acid sequence, selected the sequences with the lowest GC content and calculated the predicted melting temperature of their corresponding RNA sequence. We computed the conformations of the RNA sequences with the highest value for change in free Gibbs energy (Δ*G*) regarding the intramolecular base-pairing of the 30 bp before and 30 bp after the start codon (60 bp total) using mfold [32]. If sequences had similar Δ*G* values, the sequence with the least number of base-pairs at the ribosome binding site (RBS) and start codon was chosen. An example of the calculated RNA structures is shown in **Supplementary Fig. 1**. The sequence-optimized DNA fragments, including a T7 promotor, RBS, and T7 terminator, were synthesized and cloned in the pUC57 plasmid (GenScript, United States). Linear templates were generated by PCR using forward and reverse primers 5’-CAGTCACGACGTTGTAAAACGAC-3’ and 5’-CACACAGGAAACAGCTATGAC-3’, respectively. The PCR products were purified using the Wizard SV Gel and PCR Clean-Up System (Promega), following the manufacturer’s protocol. Concentration and purity of the DNA constructs were determined by spectrophotometry using the NanoDrop 2000 (Thermo Scientific). Samples were also analysed by electrophoresis on 1% agarose gels.

### Purification and labelling of BtubA/B

BtubA/B proteins were purified and labelled as described in [6] except that C41(DE3) cells were used for expression instead of BL21(DE3). Concentration of purified bacterial tubulin was determined from absorbance at 280 nm (extinction coefficient 103754.2 M^−1^ cm^−1^).

### Bulk CFPS

Bulk CFPS reactions were performed with PURE*frex*2.0 (GeneFrontier Corporation) and 5 nM of linear DNA template according to the supplier’s protocol. When *btubA* and *btubB* genes were co-expressed, 2.5 nM of each DNA construct was used. For visualization of gene products by PAGE, 1 µL of GreenLys solution (FluoroTect, Promega) was supplemented to a PURE*frex*2.0 mix (total volume 10 µL) and the reactions were carried out in PCR tubes at 37 °C for 3 h. Samples with added sodium dodecyl sulfate (SDS, 20% w/v final) were incubated at 90 °C for 10 min and loaded on a 12% SDS-PAGE gel that was run first for 20 min at 100 V and then for 40 min at 160 V. Fluorescently labelled proteins were visualized on a fluorescence gel imager (Typhoon, Amersham Biosciences) using a 488 nm laser and a 515-535-nm band-pass emission filter. Subsequently, the gel was stained with InstantBlue (expedeon) overnight and imaged with a ChemiDocTM imaging system (Bio-Rad).

### Estimation of the concentration of synthesized BtubA/B

Pre-ran PURE*frex*2.0 samples containing expressed bacterial tubulin were loaded onto a 10% stain-free gel, either undiluted or 5-fold diluted. Purified bacterial tubulin was loaded in concentrations ranging from 0.125 μM to 8 μM and a calibration curve was generated from these samples using Fiji [33] to quantify band intensities.

### Imaging chambers

Experiments involving SLB or liposome imaging were carried out in self-made glass chambers. Three glass slides of 1 mm thickness were glued together with NOA 61 UV-glue (Norland Products). Several holes of 3 mm diameter were created with a diamond drill and a 150-μm-thick coverslip (Menzel-Gläser) glued to the bottom with NOA 61. Before use, the chambers were cleaned by sequential washing steps of 10 min sonication each in chloroform/methanol (1:1 volume), 2% Hellmanex III (Hellma), 1 M KOH, ethanol and MilliQ water. In case of SLB experiments, the glass chambers were additionally treated every second experiment with acid piranha solution. For some liposome experiments, aluminium chambers were used, which were fabricated in the same manner as described for the glass chambers and were cleaned following the same protocol, except that the KOH and piranha treatments were omitted.

### Preparation of lipid-coated beads

All stock lipids in chloroform were supplied by Avanti Polar Lipids, except for Texas Red 1,2-dihexadecanoyl-*sn*-glycero-3-phosphoethanolamine, triethylammonium salt (DHPE-TexasRed), which was from Invitrogen. A lipid mixture consisting of ∼50 mol% 1,2-dioleoyl-*sn*-glycero-3-phosphocholine (DOPC), 36 mol% 1,2-dioleoyl-*sn*-glycero-3-phosphatidylethanolamine (DOPE), 12 mol% 1,2-dioleoyl-*sn*-glycero-3-phospho-(1’-*rac*-glycerol) (DOPG), 2 mol% 1’,3’-bis[1,2-dioleoyl-*sn*-glycero-3-phospho]-*sn*-glycerol (18:1 cardiolipin), 0.2 mol% DHPE-TexasRed and 1 mass% 1,2-distearoyl-*sn*-glycero-3-phosphoethanolamine-N-[biotinyl(polyethylene glycol)-2000] (DSPE-PEG-biotin), was prepared in a 10-mL round-bottom glass flask. A solution of 100 mM rhamnose in methanol was added (2.5:1 chloroform-to-methanol volume ratio), followed by 0.6 g of 212-300-μm glass beads (acid washed, Sigma Aldrich). The solvent was removed by rotary evaporation at 200 mbar for 2 h at room temperature. The lipid-coated beads were aliquoted, desiccated overnight and stored under argon at –20 °C.

### Preparation of SUVs

Small unilamellar vesicles (SUVs) had the same lipid composition as described for the preparation of lipid-coated beads, except that DSPE-PEG-biotin was omitted. A lipid film of 500 µg was formed at the bottom of a glass vial by gentle evaporation of chloroform. Lipids were resuspended with 400 µL MilliQ water (1.25 mg mL^−1^ final concentration) and the solution was vortexed for 2 min. Sample was extruded using a mini extruder (Avanti Polar Lipids) equipped with 250 µL Hamilton syringes, two filters (drain disc 10 mm diameter, Whatman) and a polycarbonate membrane with pore size of 0.2 μm (first extrusion) and 0.03 μm (second extrusion). The SUV stock solution was stored at –20 °C until use.

### SLB experiments

Imaging glass chambers were treated with oxygen plasma (Harrick Plasma) for 15 min to activate the surface. Six microliters of SUV solution were added to the chamber and supplemented with 12 μL of 6 mM CaCl2. The chamber was covered with a coverslip, placed on a 0.5-mm thick adhesive silicone sheet (Life Technologies), and incubated for 30 min at 37 °C. The formed SLB was washed four times with MRB80 buffer (80 mM K-Pipes, 4 mM MgCl2, 1 mM EGTA, pH 6.8) and incubated 10 min with 0.5 mg mL^−1^ k-Casein in MRB80 buffer. For experiments with purified BtubA/B, 20 μL of a solution containing PURE*frex*2.0, 0.05% (w/v) methylcellulose, 4 μL MRB80, 3 μL MilliQ water, purified unlabelled and Atto488/Atto561-labelled BtubA/B, were added to the glass chamber. Imaging was performed at either 25 or 30 °C. For experiments with cell-free synthesized BtubA/B, 8.5 μL of a pre-ran PURE*frex*2.0 sample containing expressed BtubA was mixed with 1.5 μL of 1 μM labelled BtubA/B-Atto488, the sample was added to the SLB and imaged for 30 min at 30 °C. Then, 5 μL of a pre-ran PURE*frex*2.0 sample containing expressed BtubB was added during total internal reflection fluorescence (TIRF) imaging. The setup consisted of an Ilas2 system (Roper Scientific) on a Nikon Ti-E inverted microscope with a Nikon CFI Plan Apochromat 100 × NA1.45 TIRF oil objective and two Evolve 512 EMCCD camera’s (Photometrics) for simultaneous dual-acquisition. The system was operated with MetaMorph 7.8.8.0 (Molecular Device).

### CFPS in liposomes

Twenty micrograms of lipid-coated beads were added to 20 μL of swelling solution consisting of PURE*frex*2.0, 1 μL DnaK mix (GeneFrontier Corporation), 100 nM Atto488-BtubA/B, and 3.75 nM of *btubA* and 2.5 nM of *btubB* DNA constructs. The sample was incubated on ice for 2 h during which the tube was gently manually rotated a few times. Four freeze-thaw cycles were applied by dipping the tube into liquid nitrogen, followed by thawing at room temperature. An imaging glass chamber was incubated for 5 min with a mix of bovine serum albumin (BSA) and BSA-biotin (1:1 molar ratio, 1 mg mL^−1^, Thermo Fisher Scientific), followed by incubation with Neutravidin (1 mg mL^−1^, Sigma Aldrich). Next, 4 μL of liposome sample along with 12 μL of dilution buffer (PURE*frex*2.0 Solution I and MilliQ water (7:4 volume ratio) supplemented with 83 mg L^−1^ proteinase K) were added to the imaging chamber. Fluorescence imaging was performed with a confocal microscope (A1+ from Nikon, 100× oil immersion objective) using the 488- and 561-nm laser lines for excitation of Atto488-BtubA/B and DHPE-TexasRed, respectively. Samples were incubated at 37 °C during image acquisition.

## ACKNOWLEDGEMENTS

The authors thank Vladimir Volkov for assistance with establishing bacterial microtubules in the lab and for useful discussions. Microscopy measurements were performed at the Kavli Nanolab Imaging Center Delft. This work was financially supported by the Netherlands Organization for Scientific Research (NWO/OCW) through the ‘NanoFront – Frontiers of Nanoscience’ Gravitation grant and the ‘BaSyC – Building a Synthetic Cell’ Gravitation grant (024.003.019).

## SUPPLEMENTARY INFORMATION

Supplementary Information is available for this paper: gene sequences, supplemental methods, figures, movies and a note.

