## Supplementary Information for "Shaping liposomes by cell-free expressed bacterial microtubules"

for

### SUPPLEMENTARY METHODS

#### Sequence of ***btubA*** (5’→3’, complete linear construct)

CAGTCACGACGTTGTAAAACGACGGCCAGTCGCGAAATTAATACGACTCACTATAGGGGAATTGTGAGCGGATAACAA
TTCCCCTCTAGAAATAATTTTGTTTAACTTTAAGAAGGAGATATACATATGAAAGTTAATAATACAATTGTAGTTAGTAT
TGGTCAGGCGGGCAACCAAATCGCGGCGAGCTTCTGGAAAACCGTGTGCCTGGAGCACGGTATTGACCCGCTGACCGG
TCAGACCGCGCCGGGCGTTGCGCCGCGTGGTAACTGGAGCAGCTTCTTTAGCAAGCTGGGCGAGAGCAGCAGCGGTAG
CTACGTGCCGCGTGCGATCATGGTTGATCTGGAACCGAGCGTGATTGACAACGTTAAAGCGACCAGCGGCAGCCTGTTC
AACCCGGCGAACCTGATTAGCCGTACCGAGGGCGCGGGTGGCAACTTTGCGGTTGGTTACCTGGGTGCGGGTCGTGAG
GTGCTGCCGGAAGTTATGAGCCGTCTGGATTATGAAATCGACAAGTGCGATAACGTGGGTGGCATCATTGTTCTGCATG
CGATCGGTGGTGGCACCGGCAGCGGTTTTGGCGCGCTGCTGATCGAGAGCCTGAAGGAAAAATACGGCGAGATTCCGG
TGCTGAGCTGCGCGGTTCTGCCGAGCCCGCAGGTGAGCAGCGTGGTTACCGAGCCGTATAACACCGTTTTTGCGCTGAA
CACCCTGCGTCGTAGCGCGGATGCGTGCCTGATCTTCGATAACGAAGCGCTGTTTGACCTGGCGCACCGTAAATGGAAC
ATTGAGAGCCCGACCGTGGACGATCTGAACCTGCTGATCACCGAAGCGCTGGCGGGCATTACCGCGAGCATGCGTTTCA
GCGGTTTTCTGACCGTGGAAATCACCCTGCGTGAGCTGCTGACCAACCTGGTTCCGCAACCGAGCCTGCACTTCCTGAT
GTGCGCGTTTGCGCCGCTGACCCCGCCGGATCGTAGCAAGTTCGAGGAACTGGGTATCGAGGAAATGATTAAAAGCCT
GTTCGACAACGGCAGCGTGTTTGCGGCGTGCAGCCCGATGGAAGGTCGTTTTCTGAGCACCGCGGTTCTGTATCGTGGC
ATCATGGAGGATAAACCGCTGGCGGATGCGGCGCTGGCGGCGATGCGTGAAAAGCTGCCGCTGACCTACTGGATTCCG
ACCGCGTTCAAAATTGGCTATGTTGAGCAGCCGGGTATTAGCCACCGTAAAAGCATGGTGCTGCTGGCGAACAACACC
GAAATCGCGCGTGTTCTGGATCGTATTTGCCACAACTTCGACAAGCTGTGGCAACGTAAAGCGTTTGCGAACTGGTATC
TGAACGAGGGTATGAGCGAGGAACAGATCAACGTGCTGCGTGCGAGCGCGCAAGAACTGGTGCAGAGCTATCAAGTTG
CGGAGGAAAGCGGCGCGAAGGCGAAAGTTCAAGACAGCGCGGGTGATACCGGTATGCGTGCGGCGGCGGCGGGTGTG
AGCGACGATGCGCGTGGTAGCATGAGCCTGCGTGACCTGGTTGATCGTCGTCGTTAAGCGATCACTAGCATAACCCCTT
GGGGCCTCTAAACGGGTCTTGAGGGGTTTTTTGGGCGTAATCATGGTCATAGCTGTTTCCTGTGTG

#### Sequence of ***btubB*** (5’→3’, complete linear construct)

CAGTCACGACGTTGTAAAACGACGGCCAGTCGCGAAATTAATACGACTCACTATAGGGGAATTGTGAGCGGATAACAA
TTCCCCTCTAGAAATAATTTTGTTTAACTTTAAGAAGGAGATTTGAAAATGAGAGAAATATTAAGTATACATGTAGGTC
AATGCGGCAACCAGATCGCGGATAGCTTTTGGCGTCTGGCGCTGCGTGAACACGGCCTGACCGAGGCGGGCACCCTGA
AGGAAGGTAGCAACGCGGCGGCGAACAGCAACATGGAAGTGTTCTTCCACAAGGTTCGTGACGGTAAATACGTGCCGC
GTGCGGTGCTGGTTGATCTGGAGCCGGGCGTTATCGCGCGTATTGAAGGTGGCGACATGAGCCAGCTGTTTGATGAAAG
CAGCATCGTGCGTAAAATCCCGGGTGCGGCGAACAACTGGGCGCGTGGTTATAACGTGGAGGGCGAAAAAGTTATCGA
CCAGATTATGAACGTGATCGATAGCGCGGTTGAGAAGACCAAAGGTCTGCAAGGCTTCCTGATGACCCATAGCATCGG
TGGCGGTAGCGGCAGCGGTCTGGGCAGCCTGATTCTGGAACGTCTGCGTCAGGCGTACCCGAAGAAACGTATCTTCACC
TTTAGCGTGGTTCCGAGCCCGCTGATTAGCGACAGCGCGGTGGAGCCGTATAACGCGATCCTGACCCTGCAACGTATTC
TGGACAACGCGGATGGTGCGGTTCTGCTGGACAACGAAGCGCTGTTCCGTATCGCGAAGGCGAAACTGAACCGTAGCC
CGAACTACATGGATCTGAACAACATCATTGCGCTGATTGTGAGCAGCGTTACCGCGAGCCTGCGTTTTCCGGGCAAGCT
GAACACCGATCTGAGCGAGTTCGTGACCAACCTGGTTCCGTTCCCGGGCAACCACTTTCTGACCGCGAGCTTCGCGCCG
ATGCGTGGTGCGGGTCAGGAAGGTCAAGTGCGTACCAACTTTCCGGACCTGGCGCGTGAAACCTTTGCGCAGGACAAC
TTCACCGCGGCGATCGATTGGCAGCAAGGTGTTTATCTGGCGGCGAGCGCGCTGTTCCGTGGCGATGTGAAGGCGAAA
GACGTTGATGAAAACATGGCGACCATTCGTAAGAGCCTGAACTACGCGAGCTATATGCCGGCGAGCGGCGGTCTGAAA
CTGGGCTATGCGGAAACCGCGCCGGAAGGTTTTGCGAGCAGCGGCCTGGCGCTGGTGAACCACACCGGTATCGCGGCG

GTTTTCGAGCGTCTGATCGCGCAATTCGACATTATGTTTGATAACCACGCGTACACCCACTGGTATGAAAACGCGGGTG
TTAGCCGTGACATGATGGCGAAAGCGCGTAACCAGATTGCGACCCTGGCGCAGAGCTATCGTGATGCGAGCTAAGCAA
TAACTAGCATAACCCCTTGGGGCCTCTAAACGGGTCTTGAGGGGTTTTTTGGGCGTAATCATGGTCATAGCTGTTTCCTG
TGTG

**Sequence of *btubC*** (5'→3', from T7 promotor to T7 terminator)

TAATACGACTCACTATAGGGGAATTGTGAGCGGATAACAATTCCCCTCTAGAAATAATTTTGTTTAACTTTAAGAAGGA
GATATACATATGGGCAGCAGCCATCATCATCATCATCACAGCAGCGGCCTGGTGCCGCGCGGCAGCATGGACTCCCCTC
TCGATCGTCAGCTCGCTGCCGCCTCTCGCAGTGTCGAAGAGGCGCGGAGAATGGCTTATCACGACGATTCGAAGATCGG
GTATCTGGTGGAGCAGATCAGCGTGCTGGCAGATCTGCGGCAGAAAGAGGGAGACTTCCGCAAGGCCGAGTCGCTGTA
TCGTGAGGCGCTATTCAGCGCTCAGGAGCAGCGTAAGCCGGACCCGGAGTTGCTGACGGGCATCCATTCCTTGCTGGCG
CATCTGTATGATCGCTGGGGCCGGATGGATCTCGCCTCGCAGTTTTATGAGAAAGCCCTGAAGATCGCCGAGCGAGGCG
GCATCGCCCAGAGTGATAAGGTGGCGATCATCAAAAACAATCTGGCGATGATCTTCAAGCAGCTCCGTGACTACACCC
GTGCGGAGCAGCACTACCAAGAGGCGCTGGAAATCTTCCGCAAAACGGATGGTGAATACAGCGCTCGGGTGGCTAGCG
TTTTTAACAATCTCGGGGTGCTATATTACAGCAACCTGGAGGTCGAGCAGGCGCAGGAGATGCATGAGCATGCGTTGAC
GATTCGGCAGAGCCTTTCGAATGATCAGGCGGACTCGGGGGATCTCTCACAGACCTACATCAATCTCGGCGCTGTTTAT
AAAGCGGCGGGAGATTTTCAGAAAGCTGAGGCCTGTGTGGATCGTGCTAAGAAGCTGCGGGCCAGCATGAATGGCTAC
CACCCGCAGCCGCGCCGTGCGGCGTCTTTGCTTGTCGATAAATCCCTGTGACAAAGCCCGAAAGGAAGCTGAGTTGGCT
GCTGCCACCGCTGAGCAATAACTAGCATAACCCCTTGGGGCCTCTAAACGGGTCTTGAGGGGTTTTTTG

**Vesicle flotation assay**

Liposomes were prepared according to the protocol used to generate SUVs, as described in the main text Methods, with slight modifications. The lipid film (1000 µg) was resuspended with 100 µL of a solution consisting of a 1:3 dilution of PURE*frex*2.0 with MilliQ water (10 mg $mL^{-1}$ final concentration) and vortexed for 5 min. A 20-µL PURE*frex*2.0 reaction mix containing the appropriate DNA construct was incubated for 3 h at 37 °C. The DNA construct *btubC* was designed as described in the main text Methods for the *btubA* and *btubB* templates. Twelve microliters of a liposome solution were added and the sample was incubated for 20 min at 30 °C. Next, 80 µL of a 1:1 dilution of PURE*frex*2.0 Solution-I (buffer Solution) with MilliQ water was added and the sample was spun down at 8,000 g for 2 min. After centrifugation, the lighter vesicles formed a pellet at the top of the solution. Eighty microliters of solution underneath the pellet were harvested. The pellet was subjected to a second centrifugation, another 15 µL of the bottom solution was removed and the pellet was resuspended by pipetting up and down. Of this liposome fraction, 10 µL was collected for analysis. The 80-µL solution of the bottom fraction from the first centrifugation was spun down and a 10-µL sample taken from the middle of the tube was harvested for analysis. Equal amounts of liposome and bottom fractions were loaded on a 12% SDS-PAGE gel.

### SUPPLEMENTARY NOTE

We examined the activity of cell-free expressed BtubA/B/C by conducting liposome floatation experiments (**Fig. S4A**). BtubC was produced and co-translationally labelled with GreenLys before incubation with lipid vesicles. After centrifugation, samples were collected from the liposome (upper) and bulk (lower) fractions, loaded on gel, and the BtubC protein content was analysed by fluorescence imaging. A clear enrichment of BtubC in the liposome fraction was observed, indicating binding to the lipid membrane (**Fig. S4C**). The ability of BtubC to interact with bMTs [1,2] was assessed by supplementing the solution with purified BtubA/B. In this assay, BtubC was unmodified (expression without GreenLys) and 1.2 μM of purified BtubA/B, including 100 nM of tubulin labelled with the dye Atto488, were employed. The protein mix and lipid vesicles were incubated to enable polymerization of bMTs and membrane binding. Sample was centrifuged and the partitioning of BtubA/B between the liposome and bulk fractions was analysed by gel imaging. BtubA/B was noticeably enriched in the liposome fraction (**Fig. S4B**). This result demonstrates that cell-free expressed BtubC is able to recruit BtubA/B to lipid membranes. In the absence of BtubC, purified BtubA/B was found in both the liposome and bulk fractions. It has been reported that purified BtubC interacts with polymerized BtubA/B but not with tubulin monomers or dimers [1]. Therefore, BtubC-conditional recruitment of BtubA/B to lipid vesicles would suggest formation of bMTs. Cell-free expressed, GreenLys-labelled BtubA/B incubated with lipid vesicles partitioned in both the liposome and bulk fractions (**Fig. S4C**). When all three proteins BtubA/B/C were co-expressed, enrichment of BtubA/B in the liposome fraction was visible (**Fig. S4C**), suggesting polymerization of cell-free expressed tubulin. Together, these results demonstrate that in vitro synthesized BtubC can bind to lipid membranes, where it recruits bMTs composed of cell-free expressed BtubA/B. The fact that BtubA/B was found also in the liposome fraction in the absence of BtubC suggests weak binding with membranes, this interaction being promoted by BtubC. Moreover, BtubC has been reported to stabilize BtubA/B filaments [2], which may favour the recruitment of long bMTs to lipid membranes.

### SUPPLEMENTARY FIGURES

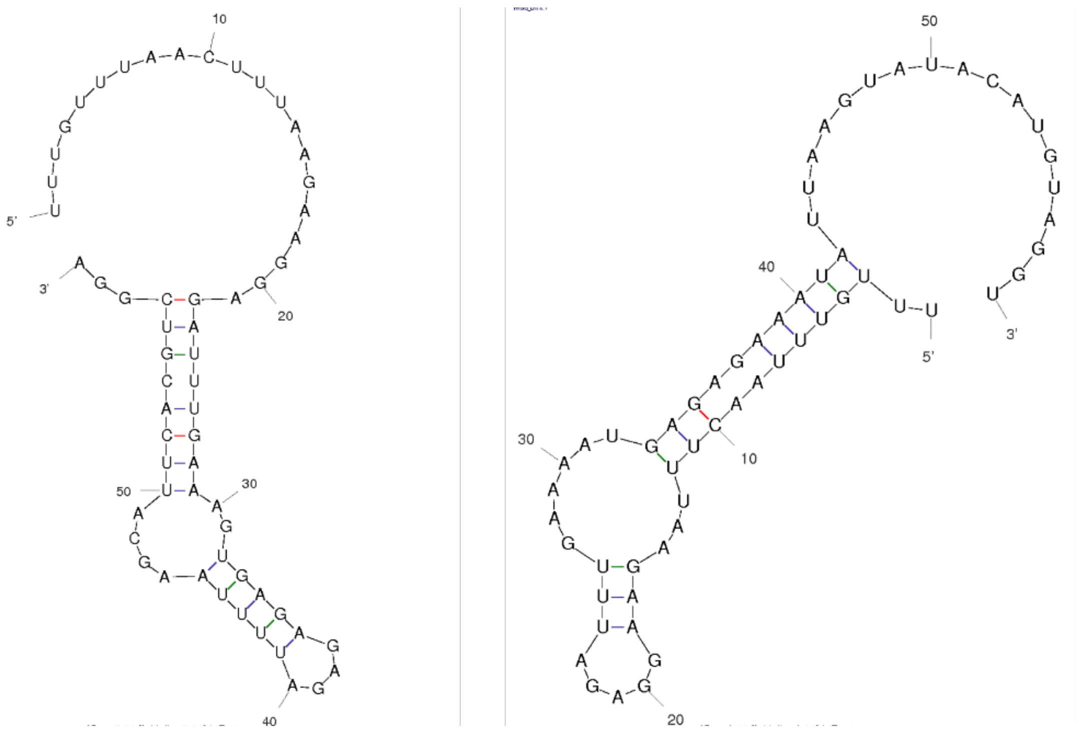


**Figure S1**: Possible RNA structures of the 30 bp sequence before and after the start codon (including start codon, 60 bp total). The structures with the lowest Δ*G* values were calculated with mfold [3] for the sequences comprising the RBS in combination with either the wildtype sequence (left image, Δ*G* = –5.8 kcal mol$^{-1}$) or the optimized sequence (right image, Δ*G* = –1.0 kcal mol$^{-1}$).

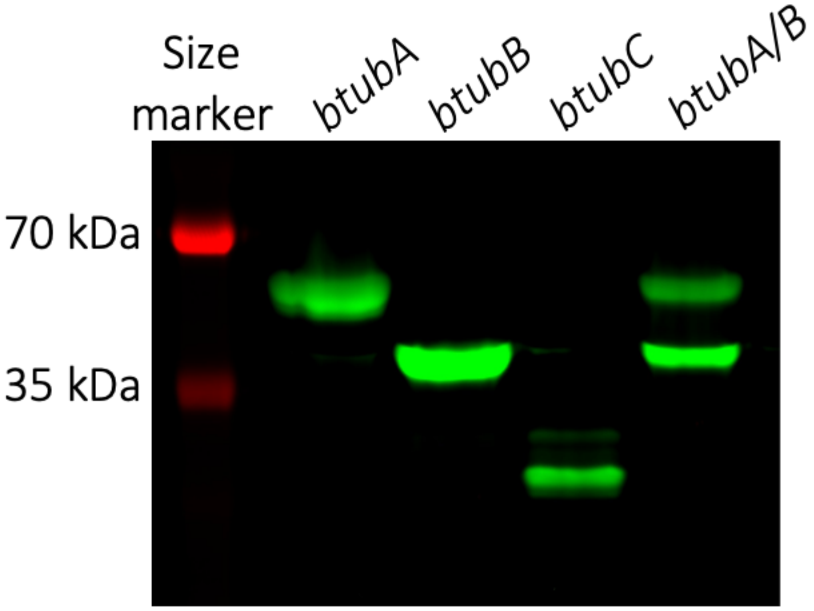


**Figure S2**: Fluorescence image of the SDS-PAGE gel shown in main text Fig. 1. Here, the lane with sample of expressed *btubC* gene is uncropped.

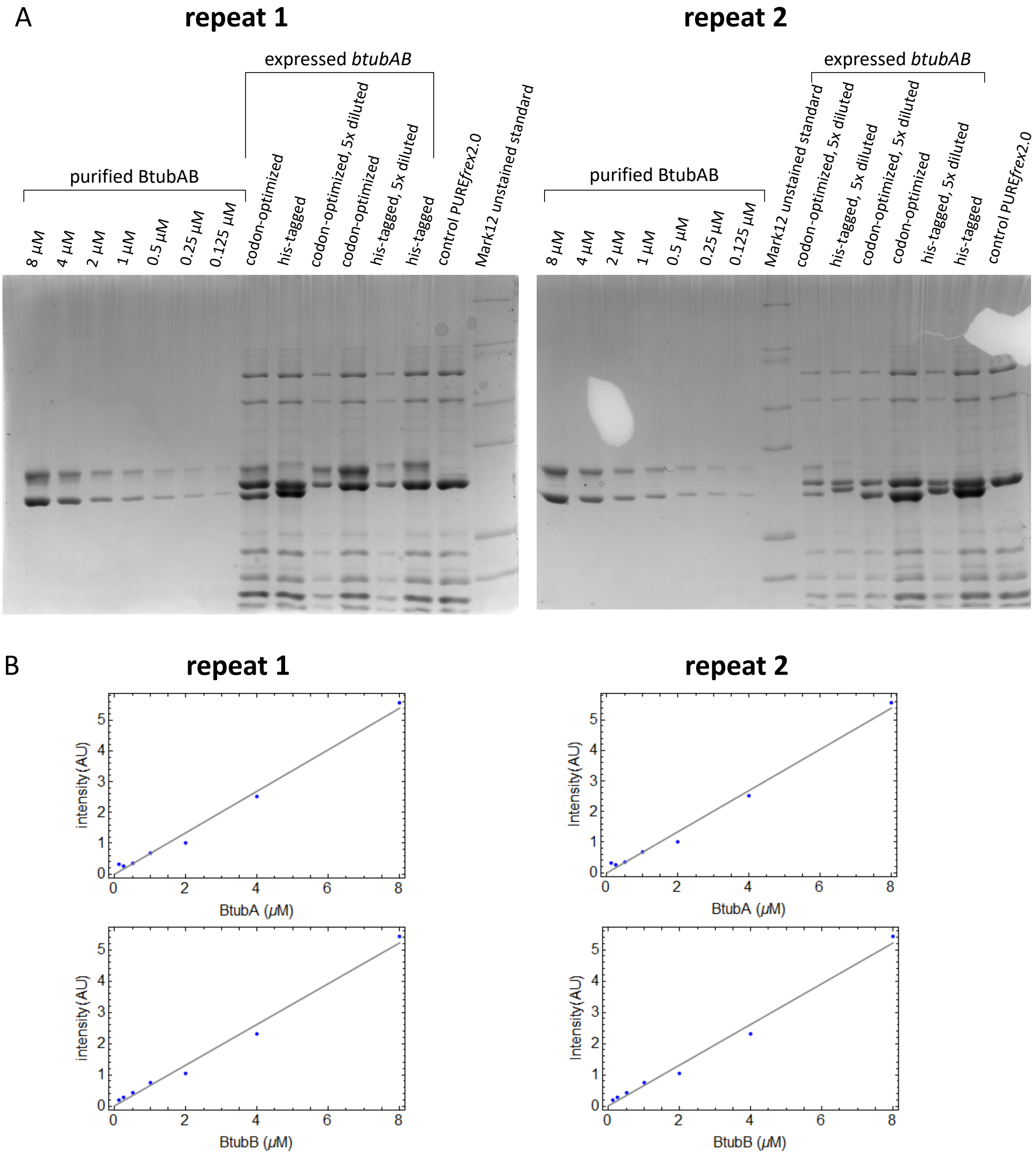


**Figure S3**: Estimation of the concentrations of expressed BtubA/B. (A) Samples of expressed and purified BtubA/B proteins were analysed by SDS-PAGE. Purified proteins were loaded at different concentrations ranging from 0.125 to 8 µM. DNA constructs of the wild-type (his-tagged) or optimized *btubA/B* gene sequences were used as templates in PURE*frex*2.0 bulk reactions. A control sample with no DNA was also prepared. Data from two independent repeats are shown. (B) Calibration curves for BtubA and BtubB were plotted by measuring the band intensity on gel for each concentration of the purified proteins. Data from the two independent experiments in (A) are shown. The measured band intensity for the 5 × diluted and undiluted PURE*frex*2.0 samples was used to extract the concentration of expressed BtubA/B.

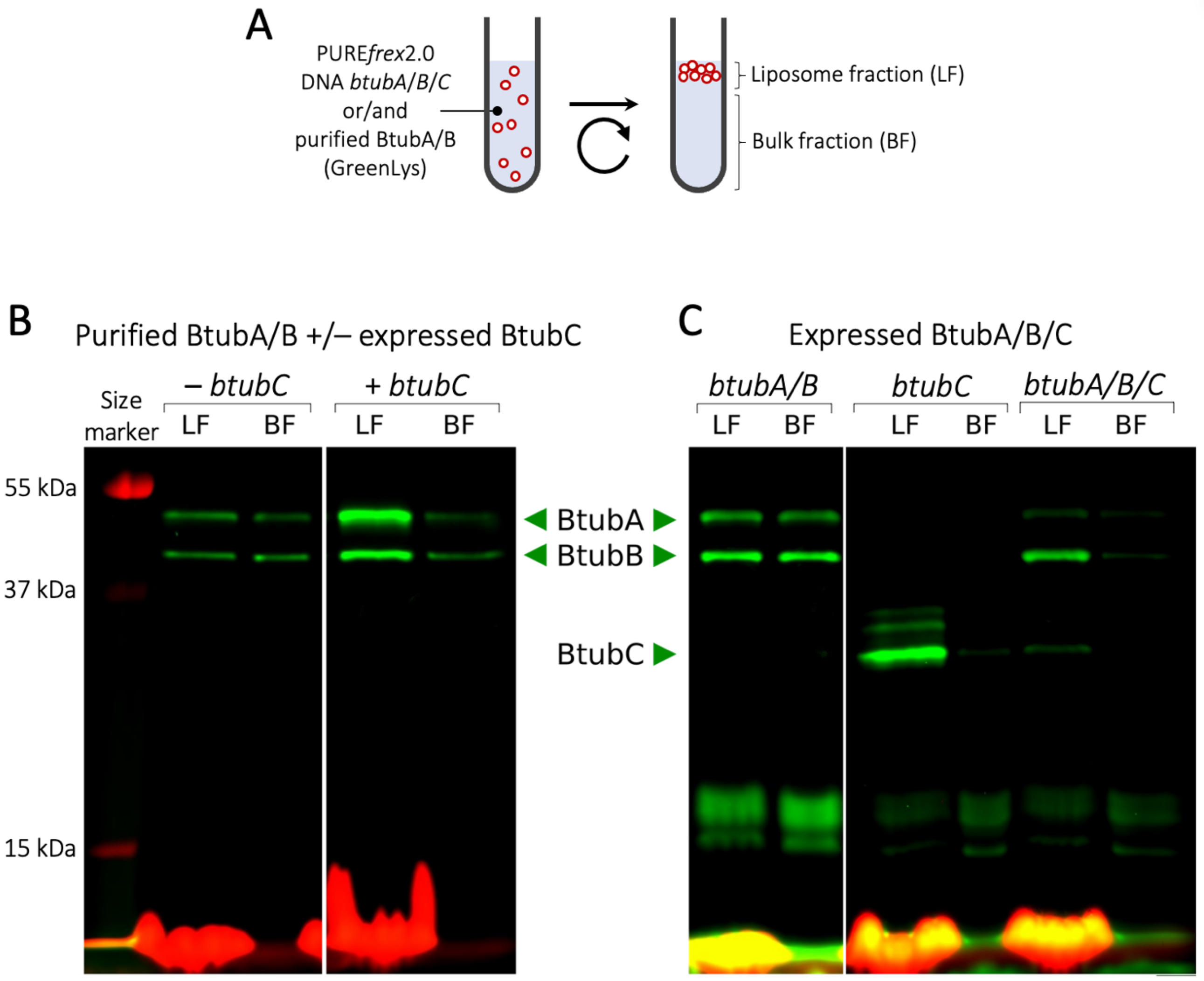


**Figure S4**: Activity of cell-free expressed BtubA/B/C characterized using a vesicle floatation assay. (A) Schematic of the vesicle flotation assay. Liposomes (coloured in red) were incubated with purified or/and pre-expressed BtubA/B/C in PURE*frex*2.0. When appropriate, the GreenLys reagent was included in the CFPS reaction. Alternatively, a fraction of purified, labelled Alexa488-BtubA/B was used for tubulin visualization by gel imaging. After centrifugation, a floating pellet of vesicles formed. Proteins interacting with the vesicles were enriched in the liposome fraction (LF) compared to the bulk fraction (BF). (B, C) SDS-PAGE analysis of BtubA/B/C partitioning between LF and BF. (B) A small amount of Alexa488-BtubA/B was used to visualize BtubAB filaments and study the ability of expressed BtubC to recruit them to the vesicle membrane. (C) BtubA/B/C were expressed with GreenLys in different combinations as indicated with the gene names. The red signal at the bottom of the lanes results from the DHPE-TexasRed lipids of the vesicles and is therefore stronger in the liposome fractions.

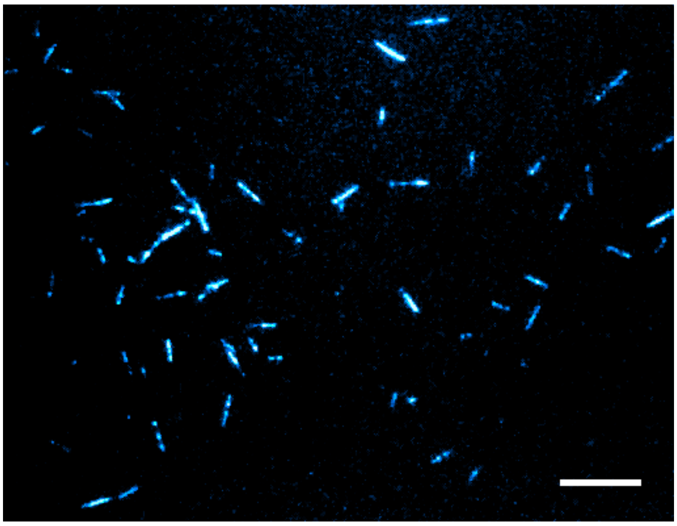

**Figure S5**: Fluorescence image of bMTs formed by 1 μM of purified BtubA/B on an SLB. Scale bar: 10 μm.

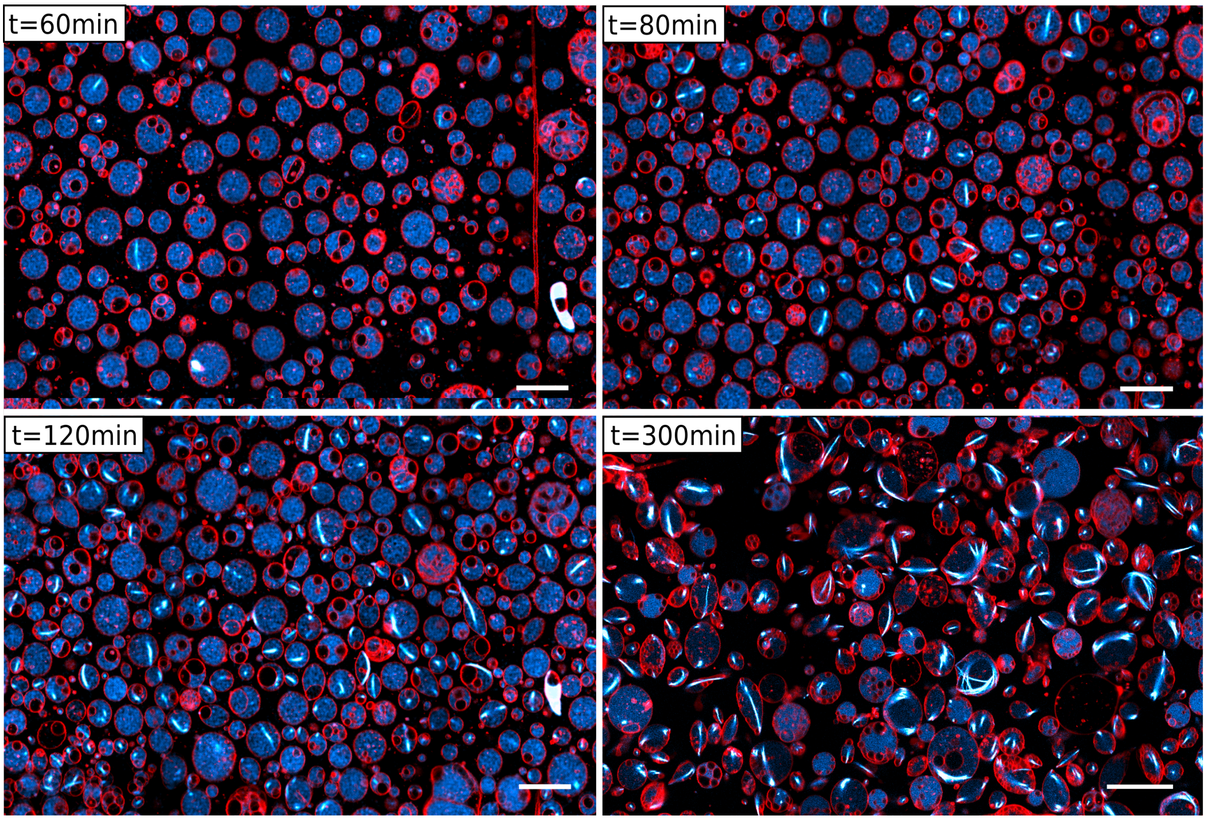


**Figure S6**: Time lapsed images of bacterial tubulin expression inside liposomes. Incubation was conducted at 37 °C. The time of incubation at which the images were acquired is appended. Scale bars: 20 µm.

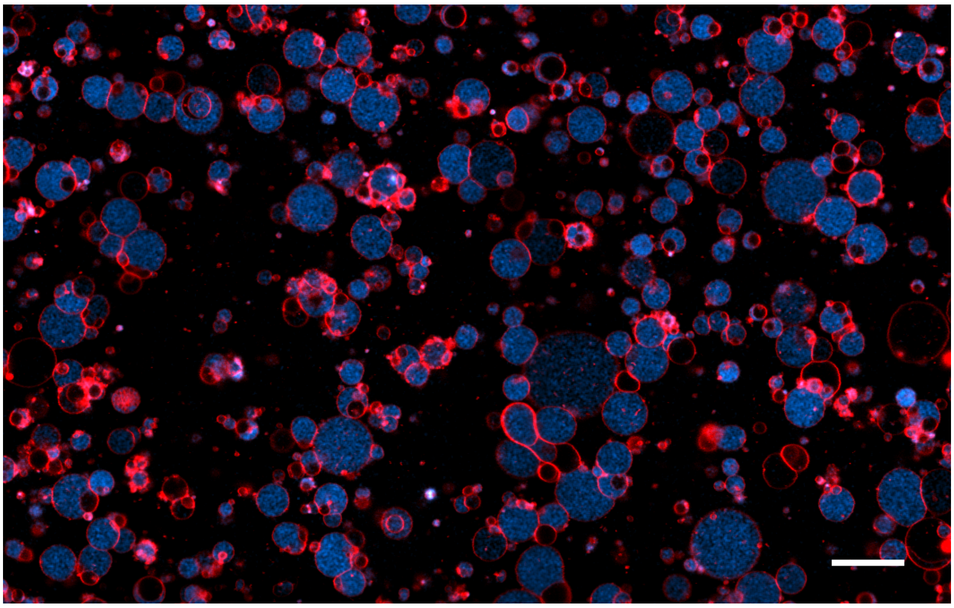

**Figure S7**: Negative control for expression of BtubA/B in liposomes. Instead of bacterial tubulin, the yeast protein Vps2 was expressed in presence of 100 nM labelled tubulin. Image acquired after 4 h incubation at 37 °C. Scale bar: 20 µm.

### SUPPLEMENTARY MOVIES

**Movie 1** TIRF imaging of purified bMTs doped with Atto488-labelled BtubA/B on a planar lipid bilayer. Experimental conditions are as described in **Fig. 2A**.

**Movie 2** TIRF imaging of purified bMTs doped with Atto488- and Atto565-labelled BtubA/B on a planar lipid bilayer. Experimental conditions are as described in **Fig. 2B-D**.

**Movie 3** TIRF imaging of cell-free expressed bMTs doped with Atto488-labelled BtubA/B on a planar lipid bilayer. Experimental conditions are as described in **Fig. 3**.

**Movie 4** TIRF imaging of purified bMTs doped with Atto488- and Atto565-labelled BtubA/B on a planar lipid bilayer exposed to intense illumination. Experimental conditions are as described in **Fig. 2E**.
